# TSG-6 in extracellular vesicles from canine mesenchymal stem/stromal is a major factor in relieving DSS-induced colitis

**DOI:** 10.1101/714931

**Authors:** Ju-Hyun An, Woo-Jin Song, Qiang Li, Min-Ok Ryu, A-Ryung Nam, Dong-Ha Bhang, Yun-Chan Jung, Hwa-Young Youn

## Abstract

Mesenchymal stem/stromal cell (MSC)-derived extracellular vesicles (EV) have been reported to be beneficial against dextran sulfate sodium (DSS)-induced colitis in mice. However, the underlying mechanisms have not been fully elucidated. We hypothesize that the tumor necrosis factor-α-stimulated gene/protein 6 (TSG-6) in EVs is a key factor influencing the alleviation of colitis symptoms. DSS-induced colitis mice (C57BL/6, male, n = 6-8/group) were intraperitoneally administered EVs (100 ug/mice) on day 1, 3, and 5; colon tissues were collected on day 10 for histopathological, qRT-PCR, western blot, and immunofluorescence analyses. In mice injected with EV, inflammation was alleviated. Indeed, EVs regulated the levels of pro- and anti-inflammatory cytokines, such as TNF-α, IL-1β, IFN-γ, IL-6, and IL-10 in inflamed colons. However, when injected with TSG-6 depleted EV, the degree of inflammatory relief was reduced. Furthermore, TSG-6 in EVs plays a key role in increasing regulatory T cells (Tregs) in the colon. In conclusion, this study shows that TSG-6 in EVs is a major factor in the relief of DSS-induced colitis, by increasing the number of Tregs in the colon.

## Introduction

Inflammatory bowel disease (IBD) is a chronic debilitating disease that affects both humans and dogs, characterized by abdominal pain and diarrhea. It may result in significant morbidity and mortality, with compromised quality of life and life expectancy. Clinical signs may be controlled by single or combination therapy, including dietary modifications, antibiotics and immune-suppressants. However, since there is no clear treatment method, clinical recurrence frequently occurs even after treatment, and thus, new therapeutic agents need to be sought [1].

Mesenchymal stem/stromal cells (MSCs) are of great interest as novel therapeutics for IBD patients because of their unique ability to regulate immune cells and heal damaged colonic tissue [2]. Particularly, it has been found that extracellular vesicles (EVs, 40–1000 nm sized circular membrane fragments shed from the cell surface) secreted from cells can mediate the delivery of secreted molecules in cell-to-cell communication, studies on the use of EVs as an alternative to stem cells have been actively conducted [3, 4]. Recently, various studies have been carried out on the application of EVs as therapeutic agents in various pre-clinical models such as acute kidney injury, hepatitis, cystitis and uveitis. Although these studies have reported that inflammation is alleviated following treatment with EVs, the factors responsible for the protective effects have yet to be elucidated [5–8]. In addition, injecting EVs into DSS-induced colitis mouse models has shown that not only does it improve activity and appetite, but it also alleviates inflammation in the colon [9].

Although the protective effects exhibited by EV has been suggested to be related to immunomodulatory factors such as transforming growth factor-beta (TGF-β), indoleamine-2,3-dioxygenase (IDO), prostaglandin-E2 (PGE2), and nitric oxide (NO)[10], the mechanisms associated with this therapeutic mechanism have yet to be examined. If stem cell-derived EVs are to be used as therapeutic agents in the future, in-depth mechanistic studies to determine which factors are most highly associated with the ability of EVs to alleviate inflammation must be conducted.

Tumor necrosis factor (TNF)-α stimulated gene/ protein 6 (TSG-6) secreted from stem cells is a major factor responsible for regulation of inflammatory responses [11–13]. Moreover, several studies have shown that TSG-6 plays important roles in attenuating DSS-induced colitis in mice by altering the composition of immune cells in the colon [12, 14]. However, studies on TSG-6 in EVs have not yet been conducted, and, thus, further studies are required.

In particular, IBD is related to an immunological imbalance in the intestinal mucosa, which is primarily associated with cells of the adaptive immune system that respond to self-antigens produced under inflammatory conditions in such patients [15, 16]. Among the intestinal immune cells, regulatory T cells (Tregs) control the balance of immune cell functions and play critical roles in self-tolerance and homeostasis in the colon [17]. However, studies on the effect of stem cell-derived EVs on colon Tregs are lacking and need to be studied.

Therefore, this study focused on elucidating the role of TSG-6 in EVs in mitigating colitis and as well as to describe potential mechanisms responsible for any protective effects observed in DSS-induced mouse models of colitis. We also investigated the effect that EVs have on Tregs within the colon and how TSG-6 in EVs affects the Treg population in inflamed colons.

## Materials and Methods

All animal experimental procedures were approved by the Institutional Animal Care and Use Committee of Seoul National University (SNU), Republic of Korea, and all protocols were in accordance with approved guidelines (SNU; protocol no. SNU-180829-2-1).

### Isolation, culture, and characterization of cAT-MSCs

The adipose tissue was obtained from a healthy adult female dog during ovariohysterectomy at the SNU Veterinary Medicine Teaching Hospital, with the owner’s consent. MSCs were isolated from tissues and cultures as previous described [12, 18]. The cells were characterized for the expression of several stem cell markers by flow cytometry before they were used in the experiments. Additionally, the differentiation ability of cells was confirmed for cAT-MSCs at passages 3 and 4, and these cells were used in subsequent experiments. The methods used for isolating, culturing, and characterizing stem cells are described in detail in supplementary material 1, and corresponding results are described in detail in supplementary figure 1. This experiment was repeated a minimum of three times to confirm reproducibility.

### Isolation and characterization of EVs

cAT-MSCs were cultured for 48 h in Dulbecco’s Modified Eagle’s Medium (DMEM; PAN-Biotech, Aidenbach, Germany) supplemented with 10 % Exosome-depleted Fetal bovine serum (FBS; Systembio, CA, USA) and 1 % penicillin-streptomycin (PS; PAN-Biotech). The supernatant from each cultured cAT-MSC sample was collected on ice and centrifuged at 300 × *g* for 10 min to remove the cells. Each supernatant was transferred to a fresh tube, centrifuged at 2000 × *g* for 30 min to remove cellular debris, and then passed through a 0.22-μm filter (Millipore, Billerica, MA, USA) to remove the large vesicles. Each supernatant was transferred to a fresh tube and centrifuged at 110,000 × *g* (Beckman Avanti Centrifuge J-26XP with 70Ti rotor, Brea, CA, USA) for 80 min, washed with Dulbecco’s phosphate-buffered saline (DPBS), and purified by centrifugation at 110,000 × *g* for 80 min. All centrifugation steps were performed at 4 °C. Each pellet was resuspended in DPBS and sterilized by filtration through a 0.22-μm filter (Figure 1C). The total protein concentration in each EV preparation was quantified by performing bicinchoninic acid (BCA) assays, and the samples were stored at −80 °C until use.

**Figure 1.**
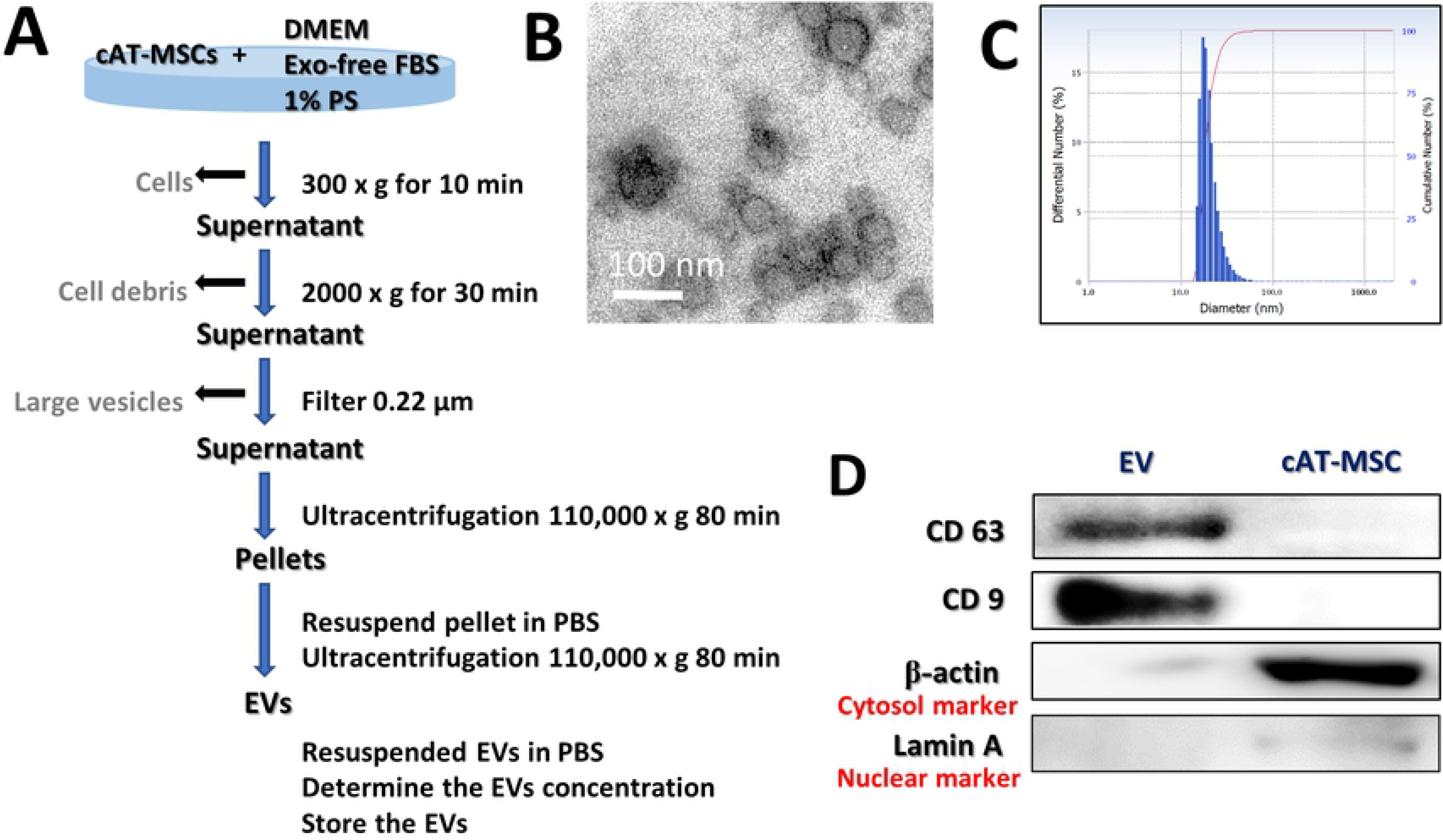
Characterization of cAT-MSC-EVs. (A) Schematic overview of the UC-based purification protocol (B) Scanning electron microscopy micrographs of cAT-MSC-EV showed spheroid shaped vesicles with diameters of approximately 20–100 nm. Scale bar, 50 nm. (C) Size-distribution analysis of purified cAT-MSC-EVs showed that the vesicle diameters were approximately 20–100 nm. (D) Immunoblotting analysis of common EV markers, where 10 μg of total protein was loaded in each lane. cAT-MSCs-EVs expressed CD63 and CD9, while beta actin and lamin A showed lower expression. The displayed data represent at least three repeated experiments with consistent results.

Protein markers of purified EVs were determined by western blotting with antibodies against cluster of differentiation 63 (CD63; NBP2-42225; Novus Inc., Littleton, CO, USA) and CD9 (GTX76185; GeneTex, Irivine, CA, USA). The EV morphology was characterized by transmission electron microscopy. Briefly, 10 μL of an EV suspension was placed on clean parafilm. A 300-mesh formvar/carbon-coated electron microscopy grid was floated on the drop, with the coated side facing the suspension, and allowed to adsorb for 20 min at 23 ± 2°C. The grid was transferred to a 100 μL drop of distilled water and left to stand for 2 min. The grid was then transferred to a 50 μL drop of 2 % uranyl acetate for negative staining for 10 min, followed by observation under a TEM (LIBRA 120, Carl Zeiss, Germany) at 120 kV. The size distribution of the particles was measured using a Zeta-potential & Particle size Analyzer (ELSZ-1000ZS, Otsuka Electronics, Osaka, Japan). This experiment was repeated a minimum of three times to confirm reproducibility.

### Isolation of peripheral blood mononuclear cells (PBMCs) and non-adherent cells

Using citrate phosphate dextrose adenine-containing tubes, we collected blood samples (30 mL) from three healthy dogs. The blood samples were diluted with an equal volume of phosphate-buffered saline (PBS) and then layered over Ficoll-Plaque PLUS (GE Healthcare Life Sciences, Little Chalfont, UK) in a conical tube. After centrifugation at 400 × *g* for 30 min, the buffy coat layer was carefully collected. The collected samples were incubated with red blood cell-lysis buffer at room temperature for 10 min. After adding PBS, each sample was centrifuged at 400 × *g* for 10 min, and the washing and centrifugation steps were repeated. Canine PBMCs (cPBMCs) were resuspended in Roswell Park Memorial Institute (RPMI) medium (Pan-Biotech, Dorset, Germany) supplemented with 10 % EV-free FBS and 1 % PS. Non-adherent cells were obtained after 24 hours.

### Treatment of cPBMC-derived lymphocytes with cAT-MSCs or cAT-MSC-derived EVs

Non-adherent cells (comprising peripheral blood lymphocytes but excluding peripheral blood monocytes) were stimulated with 5 μg/mL Concanavalin A (Con A; Sigma-Aldrich, St. Louis, MO, USA) for 12 h. The stimulated lymphocytes were transferred to a conical tube and centrifuged at 400 × *g* for 10 min. The supernatant was removed, and the remaining lymphocytes were resuspended in RPMI containing 10 % exosome-free FBS and 1 % PS. The stimulated lymphocytes (1 × 10^6^ cells/wall) were seeded in 6 well plate. Stimulated lymphocytes were cocultured with cAT-MSC-EV, siRNA-MSC-EV and siTSG6-MSC-EV (100 ug/well). Total lymphocytes without EV were cultured as a control. Total RNA was extracted from all cell groups after 48 hours of incubation.

### RNA extraction, cDNA synthesis, and reverse transcription-quantitative polymerase chain reaction

RNA was extracted using the Easy-BLUE Total RNA Extraction Kit (Intron Biotechnology). Next, cDNA was synthesized using LaboPass M-MuLV Reverse Transcriptase (Cosmogenetech, Seoul, Korea), according to the manufacturer’s instructions. cDNA samples were assayed using AMPIGENE® qPCR Green Mix Hi-ROX with SYBR Green Dye (Enzo Life Sciences, Farmingdale, NY, USA), according to the manufacturer’s instructions. Expression levels were normalized to those of glyceraldehyde 3-phosphate dehydrogenase (*GAPDH*). The sequences of the primers used in this experiment are shown in Table 1.

**Table 1.**
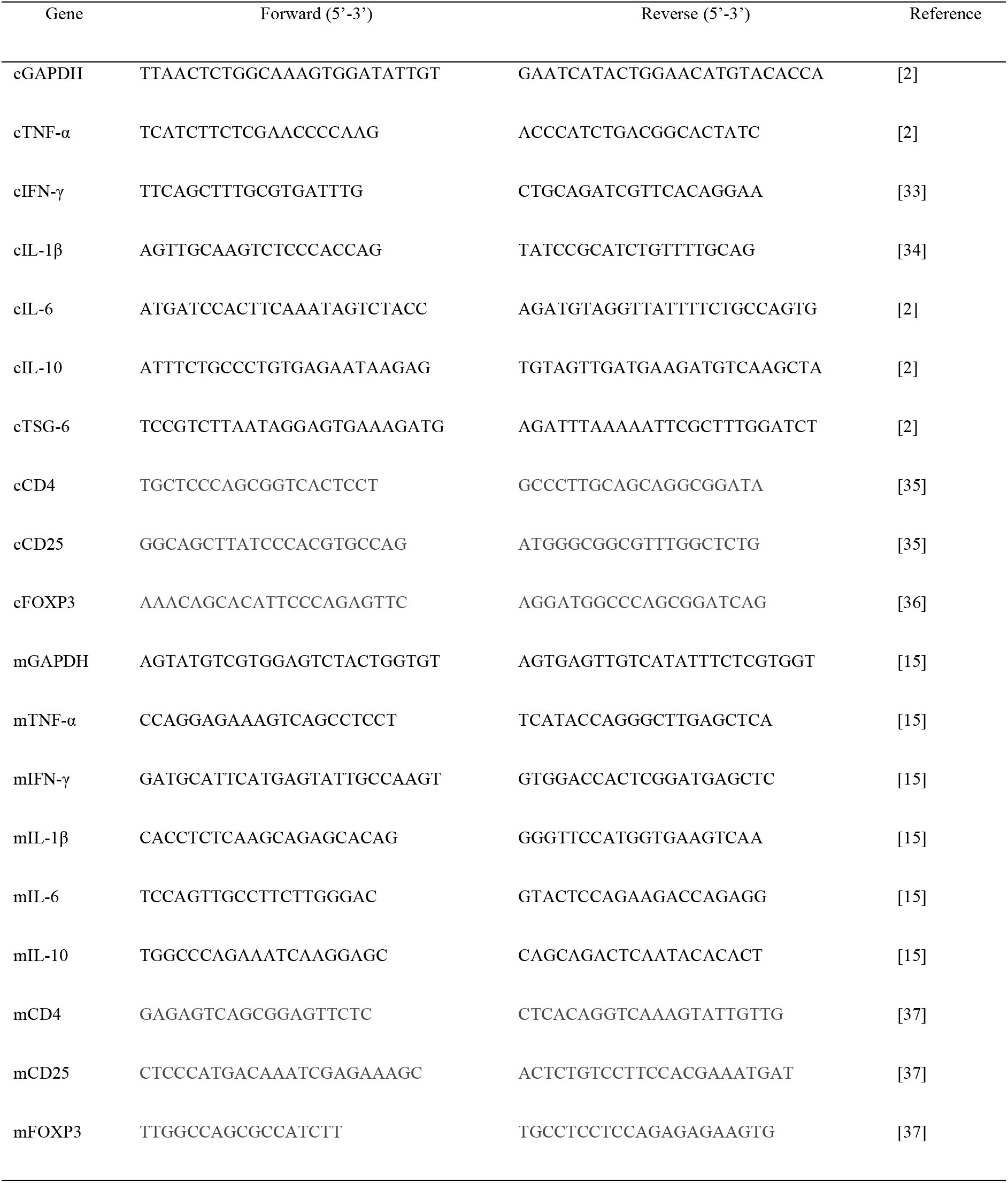
Sequences of PCR primers used in this study.

### Small interfering RNA (siRNA) transfection of cAT-MSCs

When cAT-MSCs reached approximately 70 % confluence, they were transfected for 48 h with TSG-6 siRNA or control siRNA (sc-39819 and sc-27007, respectively, Santa Cruz Biotechnology, Dallas, TX, USA) using Lipofectamine RNAiMAX (Invitrogen, Carlsbad, CA, USA) according to the manufacturer’s instructions [12, 18, 19]. TSG-6 knockdown was confirmed by qRT-PCR. The cells were washed twice with PBS, and the media was exchanged with DMEM containing Exosome-depleted FBS for an additional 48 h before collecting TSG-6-depleted EVs. EVs were obtained using an ultracentrifuge as above described, and the relative TSG-6 protein levels in EVs were measured by western blot analysis. Protein concentrations were determined by performing BCA assays. The total protein content (20 μg) of each sample was subjected to sodium dodecyl sulfate-polyacrylamide gel electrophoresis (SDS-PAGE) and immunoblotting with antibodies against CD63 (LSBio) and TSG-6 (sc-30140; Santa Cruz Biotechnology). This experiment was repeated a minimum of three times to confirm reproducibility.

### Flow cytometry analysis

To evaluate Treg polarization, PBMC-derived lymphocytes cocultured with cAT-MSC-EVs were harvested. Obtained cells (1 × 10^6)^ were suspended in 100μL DPBS and 1 μL of each primary antibody against the following proteins: FOXP3-PE (eBioscience, San Diego, CA, USA; 1:100) and CD3-FITC (MCA1774F; Bio-Rad, San Diego, CA, USA; 1:100). After incubation for 1 h at 23 ± 2 °C, the cells were washed with DPBS. Unstained cells were used as controls for autofluorescence. Cell fluorescence was analyzed with a flow cytometer (FACS Aria Ⅱ; BD bioscience). The results were analyzed using FlowJo 7.6.5 software (Tree Star, Inc., Ashland, OR, USA).

### ELISA analysis

The protein levels of interleukin 10 (IL-10) in each cell culture supernatant was detected using an IL-10 Enzyme-Linked Immunosorbent Assay Kit (ELISA; eBioscience), according to the manufacturer’s instructions.

### Mice

Male C57BL/6 mice (6-to-8 week-old and weighing 18 to 20 g) were acclimatized for 7 days with a 12 h light/dark cycle at 22 °C and 60 % humidity before performing the experiments. For environmental enrichment, 3 to 4 mice were raised in polycarbonate cages (324 × 221.5 × 130 mm) containing clean bedding (shavings; Nara Biotech), cardboard boxes, and tunnels. Mice were fed a standard laboratory rodent diet and water *ad libitum*. At the start of the experiments, the health status of the mice was evaluated by measuring their weight, vitality, and defecation; the experiments were conducted on mice with no abnormal symptoms. To generate a murine model of colitis, the mice were administered 3 % DSS (36–50 kDa; MP Biomedical, Solon, OH, USA) in their drinking water from days 0 to 7. The studies were conducted using 46 animals, and the mice were randomly divided into 6 groups with 6-8 mice per group. The mice were treated with 100 μL DPBS, with or without cAT-MSC-EV (100 μg/mouse), by intraperitoneal (IP) injection on days 1, 3 and 5. The disease-activity index (DAI) represents the combined score of weight loss relative to the initial body weight (grades 0–4; 0, no weight loss; 1, < 10 % loss; 2, 10–20 % loss; 3, 20–30 % loss; and 4, 30–40 % loss), stool consistency (grades 0–2; 0, normal; 1, soft; and 2, liquid), the presence of blood in the feces and anus (grades 0–2; 0, negative fecal occult bleeding; 1; positive fecal occult bleeding; and 3, visible fecal occult bleeding), and general activity (grades 0–2; 0, normal; 1, mildly to moderate depressed and 2, severely depressed). The DAI score of colitis was calculated independently by two blinded investigators. The score for each parameter was summed from day 0 to the day of sacrifice, and the summed score was averaged to yield the final score. On day 10 of the study, all mice were humanely euthanized with xylazine injection and CO_2_ inhalation.

### Histological evaluation

Colon tissues were fixed in 10 % formaldehyde for 48 h, embedded in paraffin, cut into 4-μm sections, and stained with hematoxylin and eosin (H&E). Histological scores are provided in Table 2. Because DSS-related injury varies, two slides from each colon section were assessed per mouse, and at least three areas on each slide were examined.

**Table 2.** Histological assessment.

| Score | Mucosal thickness & hyperplasia | Inflammatory cell extent | Damaged to the crypt |
| --- | --- | --- | --- |
| 0 | Normal | Normal | An intact crypt |
| 1 | Minimal | 11-25% of mucosa | Loss of the basal 1/3 of crypt |
| 2 | Mild | 26-50% of mucosa | Loss of basal 2/3 of crypt |
| 3 | Moderate | Mucosa and submucosa | Entire loss of crypt |

### Immunofluorescence analysis

Immunofluorescence staining was performed using mouse monoclonal anti-Forkhead box (Fox) P3 antibodies (both from Santa Cruz Biotechnology), as previously described [20]. The slides were observed using an EVOS FL microscope (Life Technologies, Carlsbad, CA, USA). Immunoreactive cells were counted in 20 random fields per section per mouse.

### Statistical analysis

Differences between more than two groups were analyzed using one-way analysis of variance (ANOVA), and the differences between two groups were analyzed using Student’s t-tests. The statistical analysis was performed using GraphPad Prism Version 6.01 (GraphPad, Inc., La Jolla, CA, USA).

## Results

### Characterization of cAT-MSC-EV

The characterization of stem cells used for these studies are described in detail in supplementary figure 1. The EVs were separated from stem cell culture media by ultracentrifugation. Schematic overview describing this protocol is presented in Figure 1A. Approximately 100 μg of EVs was produced in the media in which 1 × 10^6^ cells were seeded and grown for 2 days (data not shown). Electron microscopic analysis demonstrated that the EVs were round-shaped and 50–100 nm in diameter (Figure 1B). Using a particle-size analyzer, the EVs were confirmed to be less than 100 nm in diameter (Figure 1C). In addition, positive markers of EVs such as CD63 and CD9 were identified by western blotting, whereas negative markers of EVs such as Lamin A (a nuclear marker) and beta actin (a cytosolic marker) were present in lower abundance (Figure 1D). Whole-cell lysates were used as a positive control. Our findings suggest that the EVs contained little or no cellular matrix and nuclei, which are intracellular components.

### Production of EVs containing less TSG-6 from stem cells

To reduce TSG-6 in EVs, cAT-MSCs were transfected with si-TSG-6. No differences were observed in the cell viability and cell differentiation potentials between transfected and untreated stem cells. Furthermore, the size and shape of the EVs secreted from the transfected stem cells did not differ from those of the untreated stem cells, and no difference occurred in the amount of EVs produced (Additional Figure 1). Further, the TSG-6 mRNA levels in the transfected cAT-MSCs were reduced by over 50 % (Figure 2A), while the TSG-6 protein levels in the EVs were reduced to less than half of those in the naïve and control siRNA-treated groups (Figure 2B).

**Figure 2.**
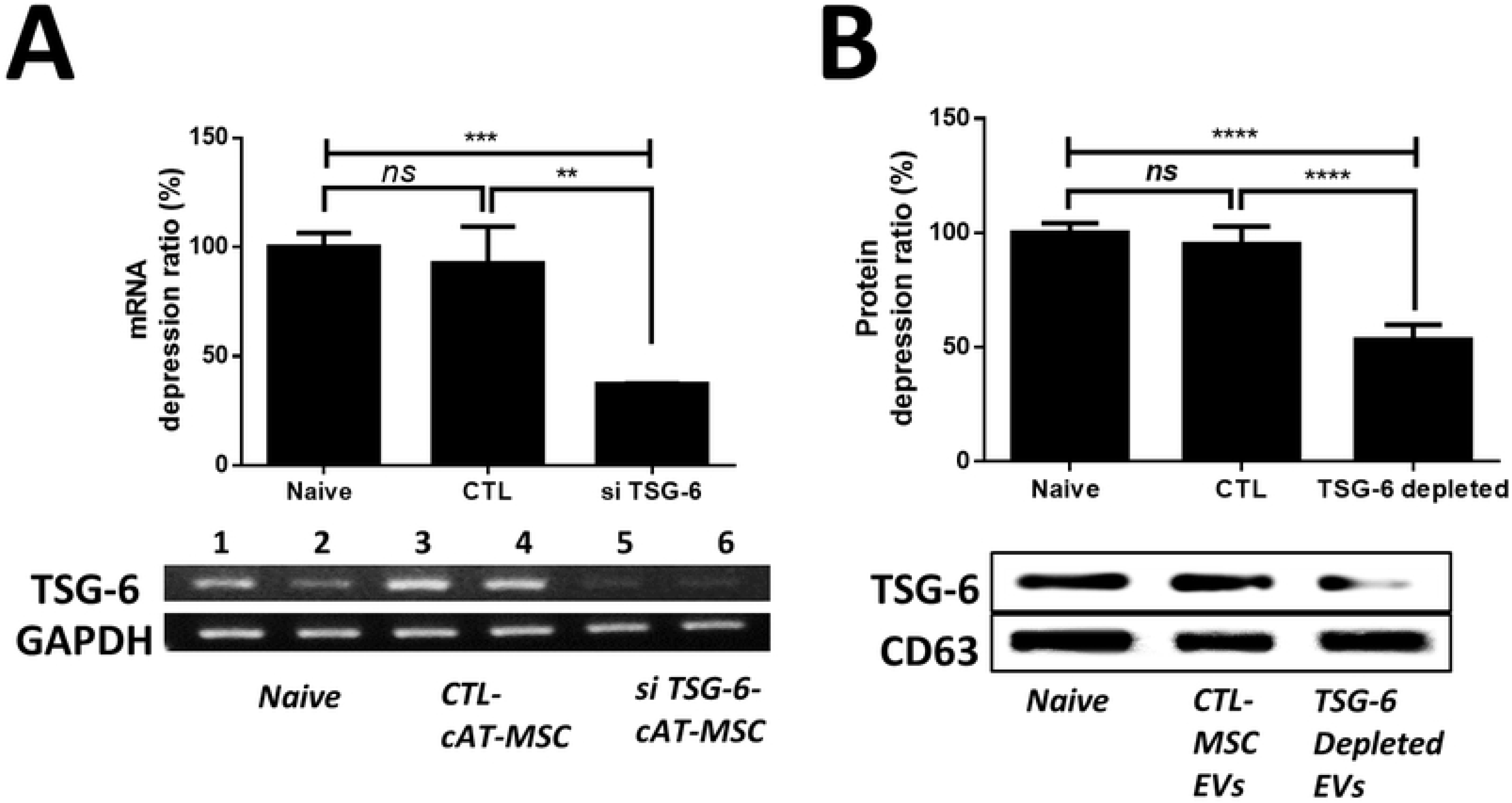
Production of TSG-6 depleted EV. (A) TSG-6 mRNA-expression levels in naïve cAT-MSCs, cAT-MSCs transfected with a scrambled siRNA (siCTL-cAT-MSC), or cAT-MSCs transfected with TSG-6 (siTSG-6-cAT-MSC) was determined by agarose gel electrophoresis and qRT-PCR. (B) TSG-6 protein-expression levels in naïve cAT-MSC-EVs, EVs from cAT-MSCs transfected with a scrambled siRNA (siCTL-EV), or EVs from cAT-MSCs transfected with TSG-6 (siTSG-6-EV) were determined by western blot analysis. The results are presented as the mean ± standard deviation. (n = 6 in each group).

### IP administration of cAT-MSC-EVs containing TSG-6 played a crucial role in alleviating IBD

Administration of EV significantly improved clinical parameters of IBD patients, such as body weight (Figure 3A). Overall, the DAI (disease activity index; considering body weight, stool consistency, bloody diarrhea and general activity) improved in the EV group (Figure 3B). Moreover, shortening of the colon length significantly improved in the EV group compared with that in the PBS-treated group (Figure 3C). The EV group showed greatly decreased histological colitis scores for mucosal thickness, mucosal hyperplasia, extent of inflammation, and crypt damage. In addition, the anti-inflammatory effect of CTL-EV was similar to that of naive-EV, whereas the effect of TSG-6-depleted EV was insignificantly decreased. (Figure 3D). Collectively, these results indicate that EV administration alleviated DSS-induced colitis and indicate that TSG-6 in EV played a role in relieving colitis.

**Figure 3.**
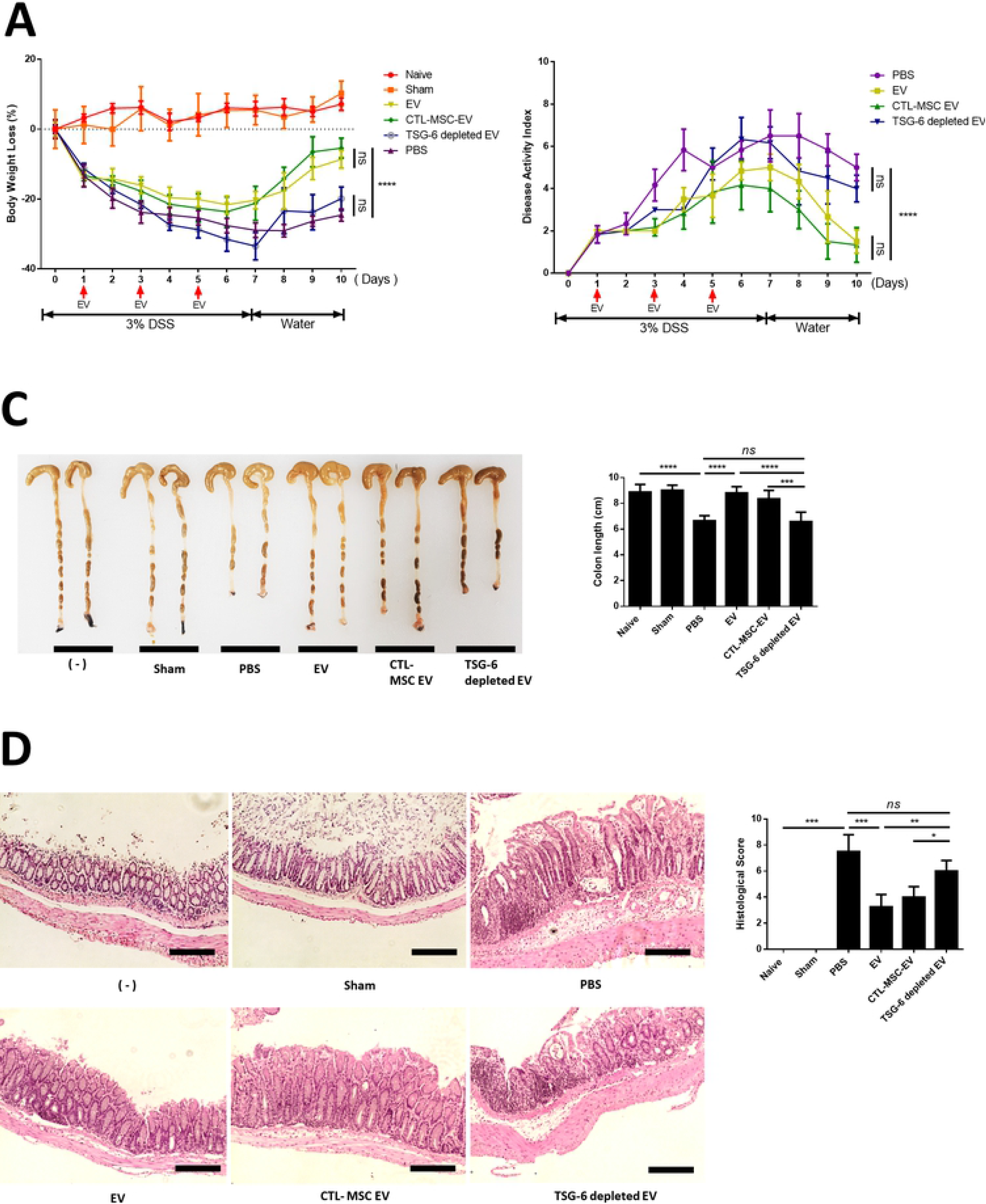
cAT-MSC-EV injection ameliorated DSS-induced colitis in mice. (A) EVs (100 μg), TSG-6 depleted EVs (100 μg), control EVs (100 μg), or vehicle control were injected IP one day after mice were administered 3 % DSS. On days 3 and 5, the mice in each group were re-injected with EVs (100 μg), TSG-6 depleted EVs (100 μg), control EVs (100 μg), or vehicle control. Mice were monitored for changes in body weight, (B) DAIs, and (C) colon lengths. (D) H&E staining of colon sections and histological scores are shown. Scale bars, 100 μm. The results are shown as mean ± standard deviation (n = 6–8 in each group, *P < 0.05, **P < 0.01, ***P < 0.001, ****P < 0.0001, as determined by one-way ANOVA). The displayed data represent at least three repeated experiments with consistent results.

### TSG-6 in cAT-MSC-EV modulate pro- and anti-inflammatory cytokine inflamed colon

The colon of EV-treated mice showed reduced levels of inflammatory cytokines (TNF-α, IFN-γ, IL-1β and IL-6) and elevated levels of an anti-inflammatory/regulatory cytokine (IL-10), compared to that of untreated mice with colitis and TSG-6-depleted EV-treated mice (Figure 4). These results indicate that EV markedly attenuated the inflammatory state in mice with induced colitis and suggest that TSG-6 in EV played an important role in relieving inflammatory conditions.

**Figure 4.**
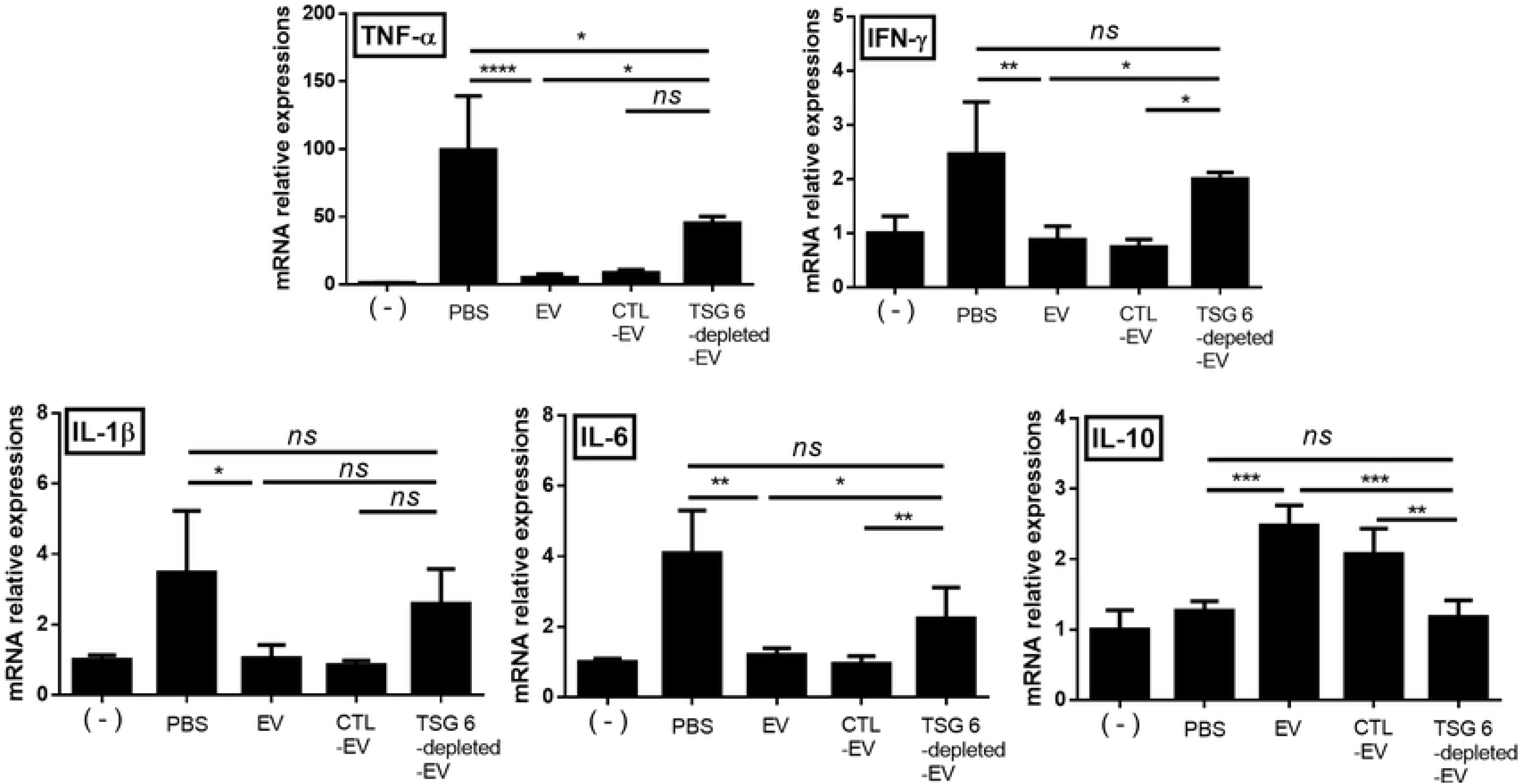
EVs from cAT-MSCs inhibited inflammatory responses in the colon. mRNA-expression levels of pro- and anti-inflammatory cytokines in the colon were determined by qRT-PCR. These data show that TSG-6 in EVs played a major role in regulating inflammatory cytokine levels in the colon. The results are shown as the mean ± standard deviation (n = 6–8 in each group, *P < 0.05, **P < 0.01, ***P < 0.001, as determined by one-way ANOVA).

### TSG-6 was a major factor in increasing Tregs in Con A stimulated lymphocytes In vitro

The expression levels of CD4, and CD25 mRNA in lymphocytes derived canine PBMCs were assessed to determine whether TSG-6 in EVs affected Treg activation. Con A, a mannose/glucose-binding lectin, is a well-known T cell mitogen that can activate the immune system, recruit lymphocytes, and elicit cytokine production. Therefore, lymphocytes were stimulated with Con A to confirm the immune cell-control function of EVs. We found that CD4, CD25 mRNA expression levels increased in the EV group, but decreased in the TSG-6 depleted EV group (Figure 5A). Additionally, the protein levels of IL-10 were measured in the lymphocyte cultured medium. The EV group showed increased IL-10 expression, whereas the TSG-6-depleted EV group showed decreased IL-10 expression (Figure 5B). In addition, to determine whether the increased number of Tregs among total T cells was associated with TSG-6 in the EVs, the degree of Treg activation was confirmed by fluorescence-activated cell sorting. Accordingly, the number of FOXP+ cells among CD3+ cells increased in the EV group. However, in the TSG-6-depleted EV group, the proportion of FOXP+ cells decreased (Figure 5C). These results demonstrate that the immunomodulatory effects of EV were related to TSG-6.

**Figure 5.**
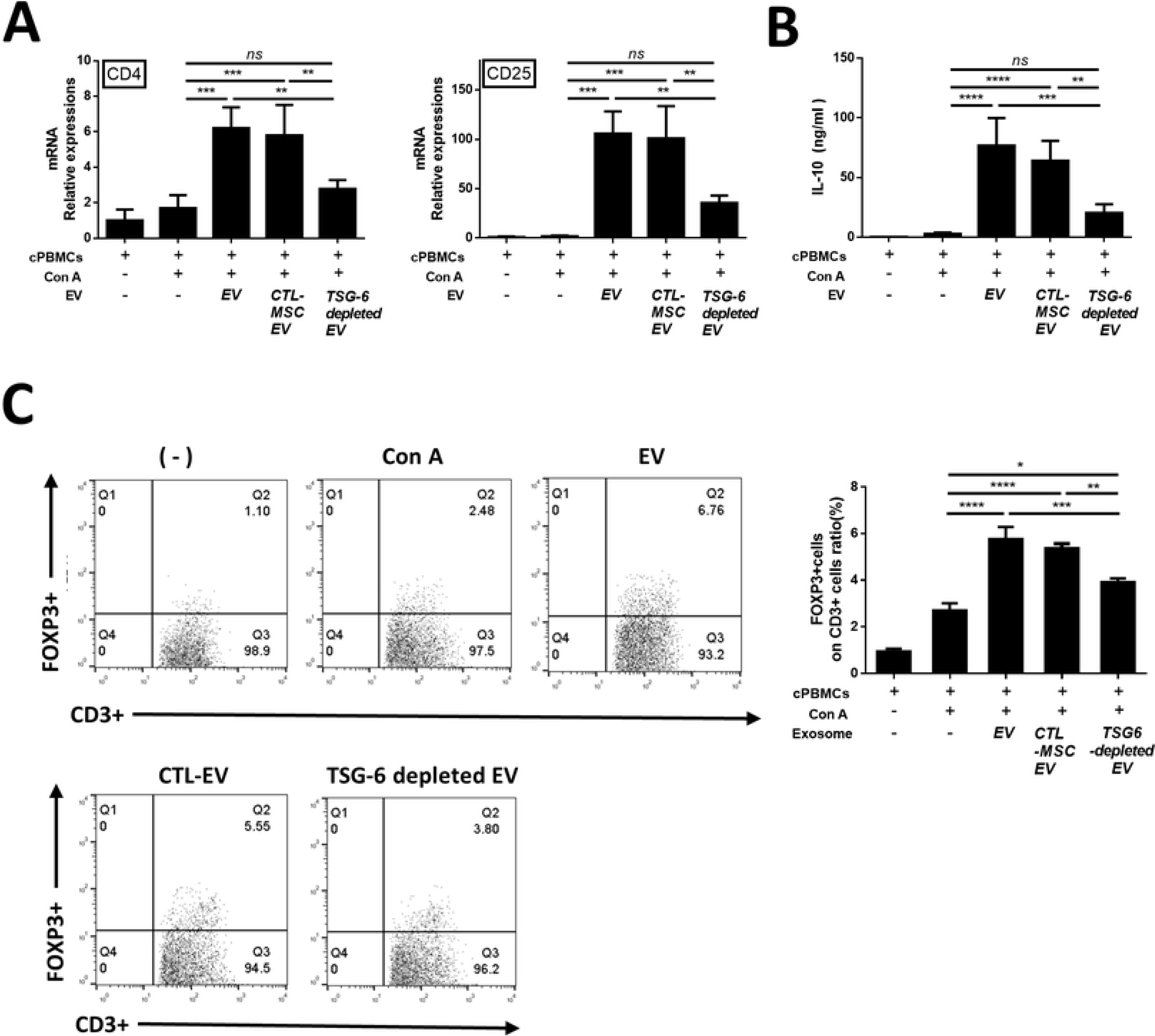
cAT-MSC-EV TSG-6 increased Treg proliferation *in vitro*. Con A-stimulated canine lymphocytes were cocultured for 48 h with cAT-MSC-EVs transfected with TSG-6 siRNA (si-TSG6) or scrambled siRNA (siCTL), or naïve EVs. (A) TSG-6 mRNA-expression levels in naïve cAT-MSCs, cAT-MSCs transfected with a scrambled siRNA (siCTL-cAT-MSC), or cAT-MSCs transfected with TSG-6 (siTSG-6-cAT-MSC) was determined by agarose gel electrophoresis and qRT-PCR. (B) TSG-6 protein-expression levels in naïve cAT-MSC-EVs, EVs from cAT-MSCs transfected with a scrambled siRNA (siCTL-EV), or EVs from cAT-MSCs transfected with TSG-6 (siTSG-6-EV) were determined by western blot analysis. The results are presented as the mean ± standard deviation of three independent experiments (n = 6 in each group). (C) CD4 and CD25 mRNA-expression levels were measured, confirming that TSG-6 was associated with increased Treg production. IL-10, which is known to be secreted from Tregs, was also measured in the supernatant medium, and the results confirmed that IL-10 production in lymphocytes was associated with TSG-6 (n = 6 in each group). (D) The Treg population was determined by measuring FOXP3 and CD3 double-positive cells by flow cytometry (n = 6 in each group). The results are presented as the mean ± standard deviation (*P < 0.05, **P < 0.01, ***P < 0.001, ****P < 0.0001, as determined by one-way ANOVA). The displayed data represent at least three repeated experiments with consistent results.

### TSG-6 in EV induced phenotypic enhancement of Tregs in inflamed colon

The FOXP3, CD4, and CD25 mRNA levels were evaluated to examine whether stem cell EVs affected the activation of Tregs in the inflamed colon. FOXP3, CD4, and CD25 mRNA levels increased in the EV group. However, their levels decreased in the TSG-6-depleted EV group (Figure 6A). To determine whether the increase in the number of Tregs was associated with TSG-6 in the EVs, quantitative analysis of FOXP3+ cells detected in colon tissue sections by immunofluorescence showed that the percentage of these cells increased significantly in the cAT-MSC-EV group compared to that in the PBS group. However, the enhancement of the number of FOXP3+ cells in the EVs of the colon tissue decreased when TSG-6 was inhibited (Figure 6B).

**Figure 6.**
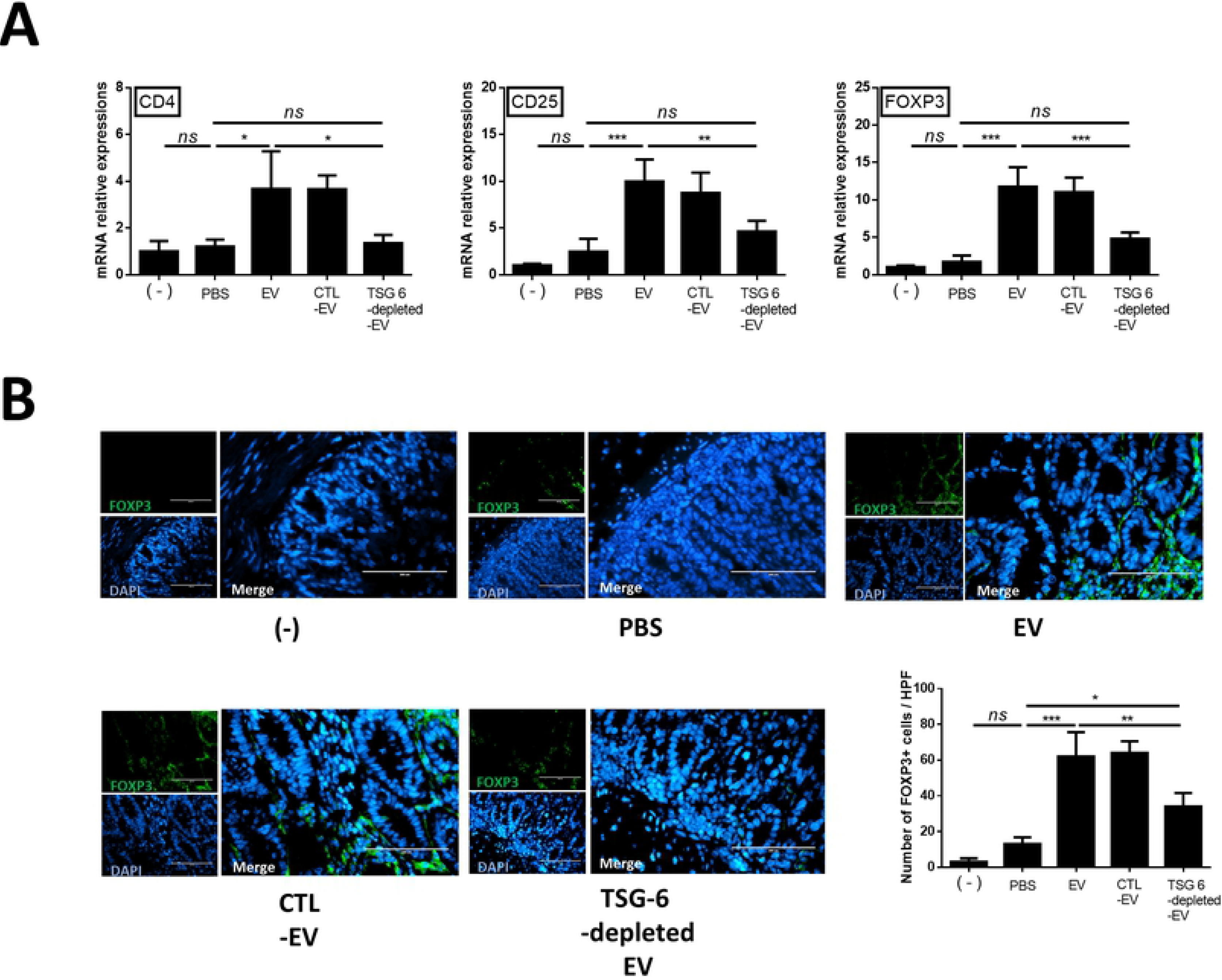
TSG-6 increases regulatory T cells in the inflamed colon. TSG-6 in EVs increased the proportion of Tregs in the inflamed colon. (A) Relative gene-expression levels of CD4, CD25, and FOXP3 in the inflamed colon. (B) Detection and quantification of FOXP3+ (green) and CD3+ (red) cells detected in colon tissue sections by immunofluorescence. The data shown demonstrated that TSG-6 in EVs played a major role in increasing the number of Tregs in the colon. Scale bar, 50 μm. The results are shown as the mean ± standard deviation (n = 6–8 in each group, *P < 0.05, ****P < 0.0001, as determined by one-way ANOVA).

## Discussion

Our data suggests the following important points: (1) TSG-6 in the stem cell derived EVs is a key factor in immune regulation and relieving inflammation in the DSS-induced mouse model of colitis. (2) TSG-6 in EVs alleviates inflammation by enhancing colonic Tregs in an IBD mouse model.

Previous studies have shown that stem cells affect recipient cells in a paracrine manner, considering that EVs largely account for the paracrine effect of stem cells [3, 21, 22]. In our, preliminary study, the immune regulatory capacity of stem cells was significantly decreased when GW4869 (a noncompetitive neutral sphingomyelinase (N-SMase) inhibitor; exosome inhibitor) was applied to cAT-MSCs (supplementary figure 2). Moreover, EVs and stem cells have similar immunomodulatory effects. These results suggested that EV can be used as a next-generation therapeutic agent as an alternative to stem cells for immune-mediated disease.

In addition, other studies have been reported to apply EVs as therapeutic agents in various inflammatory disease models and EVs have also been shown to alleviate colonic inflammation in DSS-induced IBD mouse models [3, 23]. However, although many studies have examined the therapeutic effect of EVs, the mechanism by which they relieve colitis is not yet clear. In this regard, it is noteworthy that this study demonstrated that TSG-6 in EVs is a major factor in relieving colitis symptoms.

Furthermore, this study is valuable in that it clarifies the relationship between Tregs in the colon and TSG-6 in EVs. Other studies have shown that immune cells play a role in controlling inflammation in the colon [24]. Among these immune cells, Tregs have been described as having important roles in regulating the pathogenesis of IBD [25, 26], and the balance between Tregs and other T cells in the intestinal tract is known to influence IBD pathogenesis [27].

Thus, an effective way to treat IBD may be to increase in the number of Tregs. To a large extent, FOXP3 amplifies and stabilizes the molecular features of Treg precursor cells, which is beneficial for their function and maintenance, and attenuates features that are deleterious to Treg functions [28]. FOXP3-expressing Tregs, which belong to a suppressive subset of CD4^+^ T cells, can regulate infection, tumor development, allergy, and autoimmunity [29]. It was reported that FOXP3^+^ Tregs are lower in patients with IBD progression than in healthy controls. It was also reported that an increase in the number of Tregs after treating IBD patients correlated with relief of IBD symptoms [27]. In other words, increasing the number of Tregs may serve as a method for treating IBD. In this study, we observed an increase in colonic Treg as well as a decrease in inflammatory cytokines in EV-treated IBD mice. However, treatment with TSG-6-depleted EVs did not reduce pro-inflammatory cytokines, nor did they increase the number of Tregs. These results confirmed that TSG-6 in EVs play an important role in the regulation of inflammatory cytokines and differentiation of Tregs in the inflamed colon.

Although proteins other than TSG-6, such as TGF-beta, IDO, PGE2, and NO, may also be contributing to the protective effect of EVs in relieving inflammation[10], results of the current study confirm that TSG-6-depleted EVs significantly reduce the immunoregulatory ability, which clearly indicates that TSG-6 is a major factor in immune regulation and anti-inflammatory action. Furthermore, the finding that TSG-6 in EVs plays an important role in immune regulation will serve as evidence to support increasing the level of TSG-6 in EVs as a strategy to develop EVs with enhanced immunomodulating properties.

Although EV-specific studies have not been conducted, other studies have shown that pretreatment of stem cells with TNF-alpha (TNF-α) resulted in an increase in mRNA levels of TSG-6 in stem cells as well as increased levels of TSG-6 protein in the culture medium [12, 30]. Further studies on pre-treated stem cell-derived EVs are needed.

This study is also an important basis for future transitional studies. Like human IBD, canine idiopathic IBD is a commonly observed chronic IBD that occurs spontaneously with similar multifocal etiology due to the interactions between abnormal host immune responses, and genetic and environmental factors. Histological evaluation of intestinal biopsies reveals extensive or multifocal inflammatory cell infiltration (most commonly lymphoid evolutive, eosinophilic, and neutrophilic), with simultaneous changes in mucosal structures (e.g. villous atrophy and fusion). In severe cases, intestinal protein loss, similar to in human disease, can be observed [1, 31, 32]. This study has been carried out with EVs derived from canine cells and is of great value in facilitating subsequent experiments in dogs. Therefore, evaluating the efficacy of TSG-6 in EVs conducted in this study is valuable for applications in veterinary medicine, particularly for intractable immune-mediated diseases such as IBD, however, much of these finds may also be applicable to human IBD in the future.

## Conclusion

We demonstrated that TSG-6 in EVs secreted from cAT-MSCs ameliorated DSS-induced colitis in mice by enhancing the Treg population in the inflamed colon. Our findings provide an insight to improve the current understanding of the role that EVs have in immunoregulation, and serve as a foundation for applying EVs as a therapeutic agent in IBD.

## Abbreviations

*ANOVA*: Analysis of variance
*BCA*: Bicinchoninic acid
*cAT-MSC*: Canine adipose tissue-derived mesenchymal stem/stromal cell
*CD*: Cluster of differentiation
*Con A*: Concanavalin A
*cPBMC*: Canine peripheral blood mononuclear cell
*DAI*: Disease-activity index
*DMEM*: Dulbecco’s modified Eagle’s medium
*DMSO*: Dimethyl sulfoxide
*DPBS*: Dulbecco’s phosphate-buffered saline
*DSS*: Dextran sulfate sodium
*ELISA*: Enzyme-linked immunosorbent assay
*EV*: Extracellular vesicle
*FBS*: Fetal bovine serum
*FITC*: Fluorescein isothiocyanate
*FOXP3*: Forkhead box P3
*GAPDH*: Glyceraldehyde 3-phosphate dehydrogenase
*H&E*: Hematoxylin and eosin
*IBD*: Inflammatory bowel disease
*IFN*: Interferon
*IL*: Interleukin
*IP*: Intraperitoneal
*NO*: Nitric oxide
*PBMC*: Peripheral blood mononuclear cell
*PBS*: Phosphate-buffered saline
*PE*: Phycoerythrin
*RBC*: Red blood cell
*RT-qPCR*: Reverse transcription quantitative polymerase chain reaction
*siRNA*: Small-interfering RNA
*TGF-β*: Transforming growth factor-β
*TNF-α*: Tumor necrosis factor-α
*Treg*: Regulatory T cell
TSG-6: Tumor necrosis factor-α-stimulated gene/protein-6

## Declarations

## Ethics approval

All animal experimental procedures were approved by the Institutional Animal Care and Use Committee of SNU (Protocol no. SNU-180829-2-1), Republic of Korea, and all protocols were in accordance with approved guidelines.

## Conflicts of interest

The authors declare that no conflicts of interest exist regarding the publication of this article.

## Availability of data and material

The datasets used and/or analyzed during the current study are available from the corresponding author on reasonable request.

## Funding

This study was supported by the Research Institute for Veterinary Science, Seoul National University and Basic Science Research Program of the National Research Foundation of Korea. These funds contributed to the collection, analysis, and interpretation of data generated in this study.

## Acknowledgements

Not applicable

## Authors’ contributions

JHA conceived and designed the study; collected, analyzed, and interpreted the data; and helped in writing the manuscript. WJS and QL participated in the conception and design of the study. MOR and ARN collected the data. DHB and YCJ provided administrative support and study materials. HYY contributed to the conception and design of the study, data analysis and interpretation, and granted final approval of the manuscript. All authors have read and approved the final manuscript.

## Figure legends

**Supplementary figure 1. Identification of cAT-MSCs and si TSG-6 cAT-MSC** (A) Immunophenotypes of the cultured cAT-MSCs were examined by flow cytometry. The vast majority of cells were positive for CD90, CD44, CD29, and CD73, but a few cells expressed CD34 and CD45. (B) The naïve and si-TSG-6 cAT-MSCs were maintained in specific differentiation media for 3 weeks, and the differentiated cells were stained by Oil red O to identify adipocytes, Alizarin Red S for osteocytes and Alcian Blue for chondrocytes. Scale bars, 200 μm. (C) Cell-viability assays of naïve and siTSG6-cAT-MSCs. si TSG-6 transfection was not cytotoxic when applied to stem cells. (n = 6 in each group) (D) Morphology of EVs from siTSG6-cAT-MSCs, as studied by transmission electron microscopy. EV was identified as a circular particle with a diameter of less than 100 nm. (E) EV production by naïve and siTSG6-cAT-MSCs. The production of exosome does not differ between naive and siTSG-6 groups. (n = 6 in each group) The results are shown as the mean ± standard deviation (ns, not significant, as determined by one-way ANOVA)

**Supplementary figure 2. Immunological biomarkers observed upon co-culturing total lymphocytes with cAT-MSCs** (A) Treatment with 0.005 % DMSO, 10 μM, 20 μM GW4869, or 1% DMSO showed no cytotoxic effects on cAT-MSCs, as shown by similar viability rates following all treatments, compared to the non-treated group (n = 6 in each group) (B) Pre-treatment with GW4869 (10 μM, for 12 h) significantly reduced production of EV proteins by cAT-MSCs. EV production was reduced by more than 70% in the GW4869-treated group (n = 6 in each group) (C) The mRNA levels of TNA-α, IL-1β, IL-6, IFN-γ, and IL-10 were detected by qRT-PCR. Con A-treated lymphocytes showed significantly increased levels of pro-inflammatory cytokines, such as TNF-α, IFN-γ, IL-1β, and IL-6, compared to the untreated group. cAT-MSCs depressed activated lymphocyte. however, pre-treatment with GW4869 significantly reduced the modulatory effects of cAT-MSCs. (n = 6 in each group). The results are presented as the mean ± standard deviation (**P < 0.01, ***P < 0.001, ****P < 0.0001 as determined by one-way ANOVA).

## References

1. Cerquetella M, Spaterna A, Laus F, Tesei B, Rossi G, Antonelli E, et al. Inflammatory bowel disease in the dog: differences and similarities with humans. World journal of gastroenterology: WJG. 2010;16(9):1050.

2. Yi Z, Stunz LL, Bishop GA. CD40-mediated maintenance of immune homeostasis in the adipose tissue microenvironment. Diabetes. 2014:DB_131657.

3. Katsuda T, Kosaka N, Takeshita F, Ochiya T. The therapeutic potential of mesenchymal stem cell‐derived extracellular vesicles. Proteomics. 2013;13(10-11):1637–53.

4. Burrello J, Monticone S, Gai C, Gomez Y, Kholia S, Camussi G. Stem cell-derived extracellular vesicles and immune-modulation. Frontiers in cell and developmental biology. 2016;4:83.

5. Hartjes TA, Mytnyk S, Jenster GW, van Steijn V, van Royen ME. Extracellular Vesicle Quantification and Characterization: Common Methods and Emerging Approaches. Bioengineering. 2019;6(1):7.

6. Qian X, Xu C, Fang S, Zhao P, Wang Y, Liu H, et al. Exosomal microRNAs derived from umbilical mesenchymal stem cells inhibit hepatitis C virus infection. Stem Cells Translational Medicine. 2016;5(9):1190–203.

7. Bai L, Shao H, Wang H, Zhang Z, Su C, Dong L, et al. Effects of mesenchymal stem cell-derived exosomes on experimental autoimmune uveitis. Scientific reports. 2017;7(1):4323.

8. Bruno S, Grange C, Collino F, Deregibus MC, Cantaluppi V, Biancone L, et al. Microvesicles derived from mesenchymal stem cells enhance survival in a lethal model of acute kidney injury. PloS one. 2012;7(3):e33115.

9. Mao F, Wu Y, Tang X, Kang J, Zhang B, Yan Y, et al. Exosomes derived from human umbilical cord mesenchymal stem cells relieve inflammatory bowel disease in mice. BioMed research international. 2017;2017.

10. Soleymaninejadian E, Pramanik K, Samadian E. Immunomodulatory properties of mesenchymal stem cells: cytokines and factors. American journal of reproductive immunology. 2012;67(1):1–8.

11. Lee RH, Yu JM, Foskett AM, Peltier G, Reneau JC, Bazhanov N, et al. TSG-6 as a biomarker to predict efficacy of human mesenchymal stem/progenitor cells (hMSCs) in modulating sterile inflammation in vivo. Proceedings of the National Academy of Sciences. 2014;111(47):16766–71.

12. Song W-J, Li Q, Ryu M-O, Ahn J-O, Bhang DH, Jung YC, et al. TSG-6 secreted by human adipose tissue-derived mesenchymal stem cells ameliorates DSS-induced colitis by inducing M2 macrophage polarization in mice. Scientific Reports. 2017;7(1):5187.

13. Wang N, Li Q, Zhang L, Lin H, Hu J, Li D, et al. Mesenchymal stem cells attenuate peritoneal injury through secretion of TSG-6. PloS one. 2012;7(8):e43768.

14. Milner CM, Day AJ. TSG-6: a multifunctional protein associated with inflammation. Journal of cell science. 2003;116(10):1863–73.

15. Arseneau KO, Tamagawa H, Pizarro TT, Cominelli F. Innate and adaptive immune responses related to IBD pathogenesis. Current gastroenterology reports. 2007;9(6):508–12.

16. Sartor RB. Pathogenesis and immune mechanisms of chronic inflammatory bowel diseases. American Journal of Gastroenterology. 1997;92.

17. Banz A, Peixoto A, Pontoux C, Cordier C, Rocha B, Papiernik M. A unique subpopulation of CD4+ regulatory T cells controls wasting disease, IL‐10 secretion and T cell homeostasis. European journal of immunology. 2003;33(9):2419–28.

18. Li Q, Song W-J, Ryu M-O, Nam A, An J-H, Ahn J-O, et al. TSG-6 secreted by human adipose tissue-derived mesenchymal stem cells ameliorates severe acute pancreatitis via ER stress downregulation in mice. Stem cell research & therapy. 2018;9(1):255.

19. Song W-J, Li Q, Ryu M-O, Ahn J-O, Bhang DH, Jung YC, et al. TSG-6 released from intraperitoneally injected canine adipose tissue-derived mesenchymal stem cells ameliorate inflammatory bowel disease by inducing M2 macrophage switch in mice. Stem cell research & therapy. 2018;9(1):91.

20. Ju-Hyun A, Woo-Jin S, Qiang L, Sang-Min K, Ji-In Y, Min-Ok R, et al. Prostaglandin E 2 secreted from feline adipose tissue-derived mesenchymal stem cells alleviate DSS-induced colitis by increasing regulatory T cells in mice. BMC veterinary research. 2018;14(1):354.

21. Camussi G, Deregibus MC, Cantaluppi V. Role of stem-cell-derived microvesicles in the paracrine action of stem cells. Portland Press Limited; 2013.

22. Biancone L, Bruno S, Deregibus MC, Tetta C, Camussi G. Therapeutic potential of mesenchymal stem cell-derived microvesicles. Nephrology Dialysis Transplantation. 2012;27(8):3037–42.

23. Zhang B, Yin Y, Lai RC, Tan SS, Choo ABH, Lim SK. Mesenchymal stem cells secrete immunologically active exosomes. Stem cells and development. 2013;23(11):1233–44.

24. Cook MD, Allen JM, Pence BD, Wallig MA, Gaskins HR, White BA, et al. Exercise and gut immune function: evidence of alterations in colon immune cell homeostasis and microbiome characteristics with exercise training. Immunology and cell biology. 2016;94(2):158–63.

25. Asseman C, Mauze S, Leach MW, Coffman RL, Powrie F. An essential role for interleukin 10 in the function of regulatory T cells that inhibit intestinal inflammation. Journal of Experimental Medicine. 1999;190(7):995–1004.

26. Round JL, Mazmanian SK. Inducible Foxp3+ regulatory T-cell development by a commensal bacterium of the intestinal microbiota. Proceedings of the National Academy of Sciences. 2010;107(27):12204–9.

27. Yamada A, Arakaki R, Saito M, Tsunematsu T, Kudo Y, Ishimaru N. Role of regulatory T cell in the pathogenesis of inflammatory bowel disease. World Journal of Gastroenterology. 2016;22(7):2195.

28. Tang Q, Bluestone JA. The Foxp3+ regulatory T cell: a jack of all trades, master of regulation. Nature immunology. 2008;9(3):239.

29. Sakaguchi S, Ono M, Setoguchi R, Yagi H, Hori S, Fehervari Z, et al. Foxp3+ CD25+ CD4+ natural regulatory T cells in dominant self‐tolerance and autoimmune disease. Immunological reviews. 2006;212(1):8–27.

30. Xiao Q, Wang S-k, Tian H, Xin L, Zou Z-g, Hu Y-l, et al. TNF-α increases bone marrow mesenchymal stem cell migration to ischemic tissues. Cell biochemistry and biophysics. 2012;62(3):409–14.

31. Coelho LP, Kultima JR, Costea PI, Fournier C, Pan Y, Czarnecki-Maulden G, et al. Similarity of the dog and human gut microbiomes in gene content and response to diet. Microbiome. 2018;6(1):72.

32. Vázquez-Baeza Y, Hyde ER, Suchodolski JS, Knight R. Dog and human inflammatory bowel disease rely on overlapping yet distinct dysbiosis networks. Nature microbiology. 2016;1(12):16177.

33. Schmitz S, Garden OA, Werling D, Allenspach K. Gene expression of selected signature cytokines of T cell subsets in duodenal tissues of dogs with and without inflammatory bowel disease. Veterinary immunology and immunopathology. 2012;146(1):87–91.

34. Manning K, Rachakonda PS, Rai MF, Schmidt MF. Co-expression of insulin-like growth factor-1 and interleukin-4 in an in vitro inflammatory model. Cytokine. 2010;50(3):297–305.

35. Lee K, Cha S-H, Kang H, Song J-Y, Lee K, Ko K, et al. Effects of serial passage on the characteristics and chondrogenic differentiation of canine umbilical cord matrix derived mesenchymal stem cells. Asian-Australasian journal of animal sciences. 2013;26(4):588.

36. Biller B, Elmslie R, Burnett R, Avery A, Dow S. Use of FoxP3 expression to identify regulatory T cells in healthy dogs and dogs with cancer. Veterinary immunology and immunopathology. 2007;116(1–2):69–78.

37. Jo H, Eom YW, Kim H-S, Park HJ, Kim HM, Cho M-Y. Regulatory dendritic cells induced by mesenchymal stem cells ameliorate dextran sodium sulfate-induced chronic colitis in mice. Gut and liver. 2018;12(6):664.

